# Aberrant Local Synchrony in Distinct Mouse Models of Epileptic Encephalopathy

**DOI:** 10.1101/2023.10.24.563817

**Authors:** Andrew K. Ressler, Sarah A. Dugger, Sophie Colombo, Sabrina Petri, Daniel Krizay, Wayne N. Frankel, David B. Goldstein, Michael J. Boland

## Abstract

Identifying and quantifying synchronous activity of primary neuronal networks using multielectrode arrays (MEAs) can potentially provide a medium-throughput platform to screen potential therapeutics for genetic epileptic encephalopathies (EEs). However, successfully identifying screenable synchrony phenotypes *in vitro* poses significant experimental and analytical challenges. Primary neuronal cultures quickly become highly synchronous and certain measures of synchrony tend to peak and plateau, while other network activity features remain dynamic. High levels of synchrony may confound the ability to identify reproducible phenotypes *in vitro* for a subset of EEs. Reducing, or delaying the onset of, high levels of synchrony *in vitro* may increase the dynamic range of global synchrony measures to identify disease-relevant phenotypes *in vitro,* but such measures have not been established. We hypothesized that an emphasis on local (nearby) connectivity could elucidate reproducible disease-relevant synchrony phenotypes in cortical cultures not identified by current approaches. We show clear evidence of enriched local synchrony in 48-well MEAs that varies in amplitude during development of neuronal networks. Then, we show new topological-based measures are capable of identifying novel phenotypes of aberrant synchrony in distinct mouse models of EEs. Such topological synchrony measures may provide screenable phenotypes for certain brain diseases and may be further enhanced by experimental innovation reducing global levels of synchrony in primary neuronal networks.

**Significance:** *In vitro* synchrony phenotypes may provide disease-relevant features that can be used for screening potential therapeutic candidates for epileptic encephalopathies. Here, we incorporate inter-electrode distance to generate tools capable of identifying novel synchrony phenotypes in distinct neurodevelopmental disorders. We additionally report robust topological and global *in vitro* synchrony phenotypes, alongside *in vivo* synchrony phenotypes in *Stxbp1^+/-^* mice. While singular features of disease in an *in vitro* model are unlikely to effectively test therapeutic candidates, compounds that reverse a larger subset of distinct features may translate to human patients, suggesting such a model may be ideally suited for therapeutic development using MEAs. Across multiple disease models, the topological tools developed here are complimentary to and expand upon those within meaRtools (Gelfman 2018), which is a suite of computational tools to identify network phenotypes using MEAs.

## Introduction

Aberrant neuronal synchrony is a definitive phenotype for epileptic encephalopathies; however, identifying robust synchrony phenotypes *in vitro* remains a significant challenge. Reports often do not identify specific synchrony differences or only identify them in genetically modified mouse genotypes that do not reflect those of patients (Shore 2020, Ahn 2021, Amador 2020). Further, more complex synchrony phenotypes, such as topological phenotypes that assess how groups of spatially organized neurons interact with each other can be especially difficult to identify in cultured networks. Critically, human epilepsy patients exhibit topological phenotypes. Using intracranial recordings, a breakdown of global and local topology is seen at seizure onset (Kramer 2008). Similarly, intracranial EEGs identified that slight changes in network topology could underlie the recurrence of epileptic seizures (Wang 2017). While identifying network-level aberrant synchrony either generally or topologically *in vitro* may provide a biologically relevant model to characterize and test treatments for epileptic encephalopathies, such reports remain rare.

Multielectrode arrays (MEAs) have been used to effectively study activity patterns of neuronal networks *in vitro* (see e.g., Van Pelt 2004, Esposti 2007, Massobrio 2015, Kizner 2020, Soussou 2007). Traditionally, MEA approaches utilized single wells with a high density of electrodes (Hales 2010) or require ad hoc approaches in order to record from multiple MEAs simultaneously (van Bergen, 2003). More recently, multi-well MEA systems have been developed that are capable of recording simultaneously from a number of neuronal networks, ranging from 6-well to 96-well plates. While lacking direct evidence of synchrony phenotypes in genotypes that match patients, such MEA systems have identified aberrant network activity using *in vitro* models of certain epileptic encephalopathies (‘EEs’) (Gullo 2014, Raghavan 2012, Shore 2020, Ahn 2021, Amador 2020). Interestingly, network activity of wild-type cultures shows a limited number of active electrodes (Ahn 2021) or highly synchronized activity in primary neuronal cultures (Shore 2020, Guiberson 2018, Lignani 2013) inconsistent with *in vivo* activity patterns, where networks begin to desynchronize by the end of the second postnatal week (Golshani 2009). Thus, one hypothesis for limitations of existing *in vitro* evidence is that maintained high levels of synchrony may be confounding. We set out to generate tools that assess the topology of synchrony, theorizing more complex methods with information on local networks may be needed to identify robust MEA synchrony phenotypes in models of epileptic encephalopiy in the absence of developing *in vitro* networks capable of desynchronizing similarly to *in vivo* brains at later time points.

Importantly, general topological phenotypes have been seen in cultured neuronal networks. Small-world network topology was lost in cultured neurons following the addition of glutamate (Srinivas 2007). Similarly, calcium imaging of primary neurons exposed to methamphetamine led to aberrant assortativity, despite no differences in functional connectivity (Miller 2019). Together these studies suggest topology specific phenotypes in EEs can be identified *in vitro;* however, topological approaches often focus on singular networks and are not generally designed to compare mutant and wild-type cultures across a large number of networks (Gerhard 2011, Schroeter 2015, Pastore 2021). While several computational tools exist to quantify proxies for synchrony in wild-type and mutant networks (e.g., Gelfman 2018, Eisenman 2015, Lama 2018, Bradley 2018), none specifically considers how topology, as it pertains to distance, relates to synchrony. Here, we develop new computational approaches to identity synchrony differences in primary networks from mouse models of neurodevelopmental disorders.

We first show that local electrodes have more synchronous firing than more distant electrode pairs using two new distinct measurements of local topology, the Local Synchrony Coefficient (‘LSC’) and the Local Burst Synchrony Coefficient (‘LBSC’). These measures extend two previously described quantifications of synchrony, the Spike-Time Tiling Coefficient (‘STTC’) and Mutual Information, by generating a plate-level statistic that quantifies the relationship between those synchrony features and electrode distance. We then show LSC and LBSC are capable of identifying novel phenotypes independent of global synchrony phenotypes in the presence of GABAergic and glutamatergic modulators, as well as in models of three distinct neurodevelopmental diseases. Further, we adapted a commonly used topological approach, the clustering coefficient, to identify discordant synchrony phenotypes in mice heterozygous and homozygous for a gain-of-function mutation in the NMDAR subunit, *Grin2a,* early during network development.

## Materials and Methods

Below, we outline the experimental MEA paradigm used to generate all data herein, describe the existing synchrony measures and the topological methods used to assess the networks and finally we describe the drug-dosing paradigm and genetic mouse models used to validate the potential of such analyses.

### Mice, Husbandry and Genotyping

All data were generated using mice bred and maintained at [Author University] under IACUC approval and all mouse work was in accordance with state and federal Animal Welfare Acts and Public Health Services policies. Mice were housed in ventilated cages at controlled temperature (22–23°C), humidity (∼60%), and 12h:12h light:dark cycles. Mice ate regular chow and water without restriction.

Cortical networks from *Grin2a*^S644G^, described in Amador 2020, were re-analyzed. Networks from *Hnrnpu*^+/113DEL^ mutant mice, which were generated on the C57BL/6NJ background and characterized previously (Dugger 2020), were assessed. Stxbp1 targeted mice (C57BL/6N-Stxbp1tm1a(EUCOMM)Hmgu/JMmucd) were obtained from The Jackson Labs and maintained on the C57BL/6NJ background. Cortical networks from *Stxbp1*^+/-^ and wild-type littermate controls were assessed on the F1 hybrid background B6NJ/FVB.

*Grin2a*^S644G^ and *Hnrnpu*^+/113DEL^ mutant mice were genotyped as previously described (Amador 2020, Dugger 2020). *Stxbp1*^+/-^ mice were genotyped using allele-specific PCR amplification resulting in a 444bp wild-type band and 300bp mutant band (WT F: CAA ACT CAG ACT GGC CTA TGA, WT R: AGT AAA GCA GGC AAA GAA TGC, MUT F: CGG TCG CTA CCA TTA CCA GT, MUT R: TGC AAA GAG AGG TTG GTT TGA).

### MEA Experimental Design and Statistical Analyses

#### Primary Neuron Culture

For all MEA experiments, cortical neuron dissociation, plating and network maintenance were performed similarly. Following genotyping, cortices from wild-type and mutant littermates were first separated from the rest of P0 mouse brains in Hibernate A medium (Thermo Fisher). Cortices were then dissociated into single cells via incubation in 20 U/mL Papain/DNase (Worthington) for 15-20 minutes at 37°C. Following manual trituration using a P1000 tip, cells were centrifuged at 300 x g for 5 minutes and washed in 1X PBS. Pellets were then resuspended in Neurobasal-A (Life Technologies) supplemented with 1% B27 (Life Technologies) or 1% B27 Plus (Life Technologies), 1% GlutaMax (Life Technologies), 1% HEPES, 1% Penicillin/Streptomycin, 1% fetal bovine serum (Gibco) and 5 µg/mL laminin. Live cells were counted using trypan blue and fifty thousand cells were plated in a 40-50 µl convex droplet onto wells of a 48-well MEA plate (Axion BioSystems) that had been pre-coated with 50 µg/mL poly-D-lysine (Sigma) in either 0.1M borate buffer or H_2_0, rinsed with PBS and dried for at least 1h. The following day, 500 µl fresh media was added as described above, without laminin or fetal bovine serum.

For all experiments, a 50% media change with Neurobasal-A, B27 or B27 Plus, GlutaMax, HEPES and P/S occurred every two days. Chronic drug dosing experiments were performed by adding the specified concentration of drug based on an approximation of 500 ul of media per well. Drug dosing was performed following 50% media change on DIV 3 and 5 and compounds were serially diluted with 50% media changes for the remainder of experiments.

Cultures were allowed to equilibrate for 10 min at 37°C in a carbogen mixture of 95% O_2_, 5% CO_2_ before activity recordings. Recordings were for 10 min and were always performed immediately prior to media changes using Axion BioSystems Maestro 768 channel amplifier and Axion Integrated Studios (AxIS) software (v2.4). A Butterworth bandpass filter (200-3000 Hz) and 7X standard deviation adaptive threshold spike detector were used to generate raw data and spike files.

#### Global Synchrony

Following generation of raw data using AxIS software, we used meaRtools (Gelfman 2018) to analyze a wide variety of features, including number of active electrodes, mean firing rate, mutual information and spike-time tiling coefficient (STTC). The topological analyses within this work incorporate spatial considerations to a subset of measures of synchrony from the meaRtools pipeline. Importantly, STTC and Mutual Information assess synchrony through distinct algorithms. While both algorithms rely on pairwise correlations between electrodes, the mathematical approaches are quite different. Broadly, STTC quantifies the proportion of spikes in electrode B that fall within a specified time window of a spike in electrode A and uses this as a proxy for correlation of activity in two electrodes (Cutts 2014). Alternatively, Mutual Information only considers portions of the spike-train with >75^th^ percentile levels of activity and considers these bins of activity ‘bursts.’ Then, the algorithm generates a proxy for correlated activity by giving each bin of a time a value of 1 if both electrodes are simultaneously bursting or not bursting and a 0 otherwise. While both measures are a measure of synchronous firing, they are distinct and may diverge in direction in certain disease models.

#### Local Synchrony Coefficient

Local synchrony coefficient is a topological measure that quantifies how dependent the strength of pairwise STTCs is on the distance between two electrodes. First, all pairwise STTCs of active electrodes are generated. Then, a histogram is generated with the STTCs of all active electrode pairs across all wells within a plate are binned by distance between the electrodes. For each plate, a linear regression line is fit to the histogram and the slope of the linear regression line is extracted. Finally, the local synchrony coefficient ‘LSC’ is generated by multiplying the slope by −10,000. More simply, a value greater than 0 means that there is greater synchrony between electrodes that are spatially closer than between electrodes that are further apart, suggesting nearby neurons have more tightly coupled activity patterns.

#### Local Clustering Coefficient

Clustering coefficients are a commonly used method within graph theory to measure the cohesion of a theoretical network (Chalancon 2013). More recently, clustering coefficients have been used to assess network dysfunction in epileptic patients (Pedersen 2015, Paldino 2017). Further, in high-density cultured neurons in single well MEAs, a form of clustering coefficient showed alterations in topology as the networks matured (Downes 2012). Here, we integrate the frameworks of clustering coefficient and STTCs to generate A local clustering coefficient (‘LCC’).

Broadly, a clustering coefficient is the proportion of nodes in a graph that are considered ‘connected.’ For the LCC, we restrict our analyses to local electrodes, defined by horizontal, vertical and diagonal neighbors (200um, 200um, 282.84um). For each electrode, the pairwise STTCs to all local electrodes are considered and the two electrodes are considered connected if the pairwise STTC surpasses a threshold of 0.5. Then, for each electrode a clustering coefficient is determined by the proportion of local electrodes that are connected. For each well, a single LCC is generating by averaging the LCC of all active electrodes. For this measure, the distribution of LCCs across wells is considered for downstream analyses.

#### Local Burst Synchrony Coefficient

The Local Burst Synchrony Coefficient (‘LBSC’) is similar to the LSC in determining the difference in strength of connectivity between local and more distant electrodes but the LBSC leverages the Mutual Information algorithm from meaRtools that quantifies the correlation of bursting activity between electrodes. Specifically, let LOCAL equal the aggregated pairwise Mutual Information statistics of all sets of local electrodes of a given well. Further, let DISTANT equal aggregated pairwise Mutual Information statistics of each electrode to all other electrodes in the well, excluding their local networks. We then calculate the LBSC simply with a percent increase calculation. LBSC = (local – distant / local) * 100. As with the LSC, we generate a plate level value and downstream analyses are performed with a single value per plate.

#### Statistical Analyses

The LSC and LBSC were first used to determine whether or not a topological approach to synchrony *in vitro* was applicable in 48-well MEA plates with just 16 electrodes per well. To determine the presence or absence of enriched synchrony in local networks, we assessed 220 wild-type networks (i.e. wells) across 10 litters. For both LSC and LBSC, we generated a single plate level statistic for each of the 10 litters and then used a one-sided Mann-Whitney p-value to test whether LSC and LBSC reject the null hypothesis of no enrichment of local synchrony. For comparisons between wild-type and mutant or drug dosed conditions, we used a two-sided Mann-Whitney test.

Unlike the LSC and LBSC, the LCC does not utilize a single plate-level statistic. Thus, statistical testing instead relied on a comparison of distributions. Specifically, a single plate level p-value was generated using an Earth Mover’s Distance (Urbanek 2015) calculation between wild-type and mutant or drug dosed conditions. Then, a combined p-value was generated using Stouffer’s combined method. When comparing significance of LCC to time-matched STTC, an identical one-sided Stouffer’s combined p-value was used. For all combined p-values, singular p-values were constrained with a minimum of .0001 or maximum of .9999.

#### Comparison to Non-Topological Synchrony Features

Importantly, all topological analyses described are extensions of existing measures of synchrony, STTC and Mutual Information. Thus, when determining the existence of topological phenotypes in models of disease or upon activity modulation, we similarly assessed whether or not mean firing rate, mean STTC or Mutual Information more generally showed correlated disruption. To this end, we used meaRtools to generate all of these features as previously described and available on CRAN (https://cran.r-project.org/web/packages/meaRtools/index.html). When compared with LSC and LBSC, the distributions of plate-matched mean STTC and mutual information were extracted and p-values were identically generated using a two-sided Mann-Whitney test.

#### EEG of Stxbp1^+/−^ mice

Electrodes were implanted in 8- to 10-week-old F1 hybrid wild-type and *Stxbp1*^+/−^ mice, as previously described (Asinof 2015). Briefly, three silver wire electrodes were placed 1 mm rostral to bregma and 1 mm to either side of midline subdurally, with a fourth electrode over the cerebellum as reference. An 8 research amplifier (Natus, Inc) was used to acquire signal in 48-hr continuous recordings. NeuroWorks software (Natus, Inc) was used to examine referential montages. Spike-and-wave discharges (‘SWDs’) were scored manually. Events shorter than 1s, or with amplitudes less than twice the adjacent background, were excluded from analysis.

## Results

For initial characterization of presence or absence of enriched synchrony among local electrodes, we considered 10 litters of wild-type networks with recordings from eleven days *in vitro* (DIV11) through DIV21. Further, we ensured all plates and DIVs considered had at least 11 active electrodes at all time points. Raster plots show widespread activity across electrodes, with increasing levels of activity over time, which can be visualized by representative raster plots for DIVs 11,15 and 19 shown in figure 1a.

**Figure 1:**
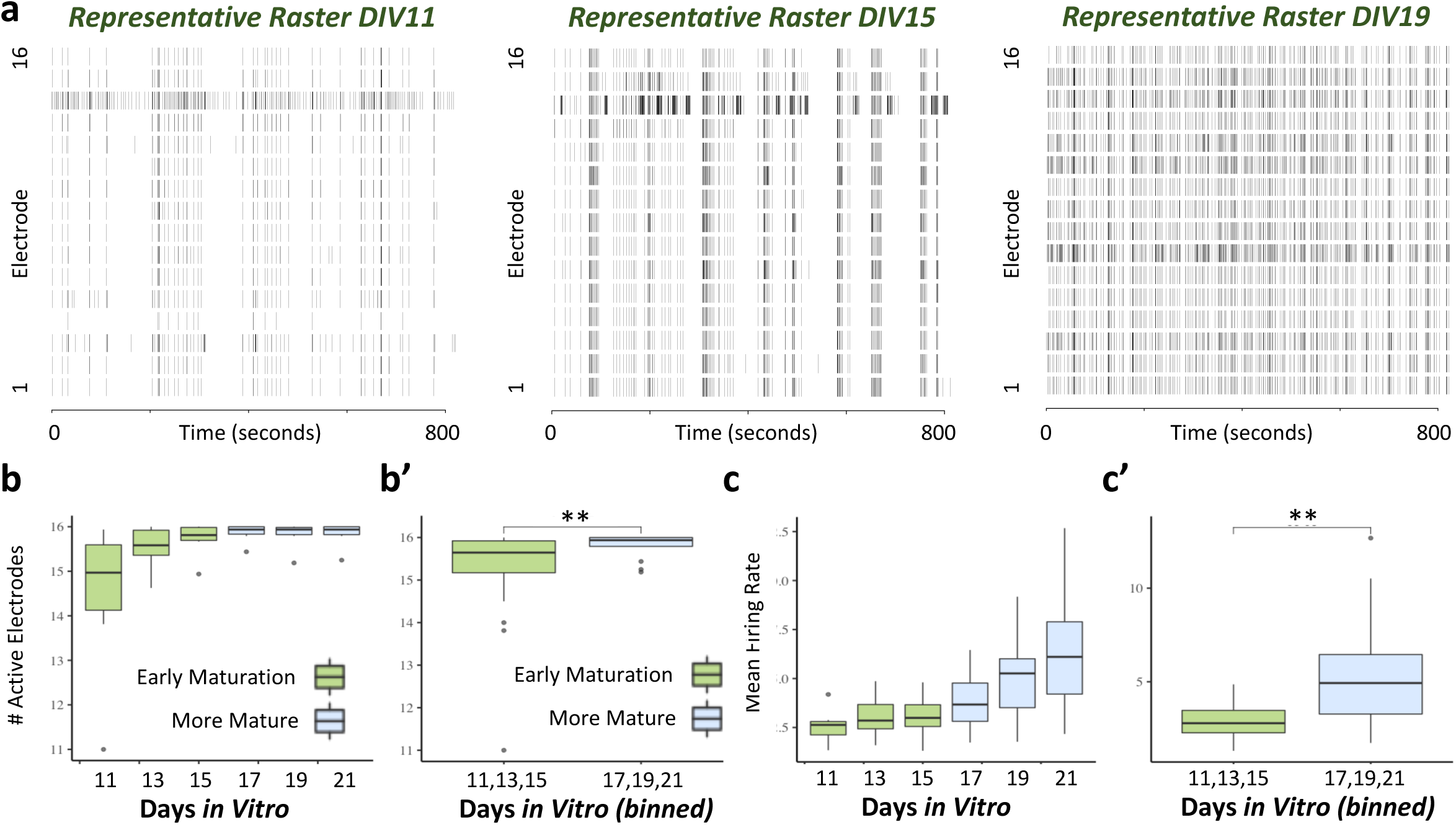
Topology in MEAs. a) Schematic of meaRtools calculation of STTC. For each electrode, an STTC value is generated by averaging the STTC of all active electrode pairs. a’) Calculation of local synchrony coefficient (LSC). LSC is calculated by generating pairwise STTCs for all electrode pairs and regressing based on distance. a’’) LSC shows enriched local topology emerging at DIV13. b) Schematic of meaRtools calculation of mutual information (MI). For each electrode, a singular value is calculated by averaging pairwise MI of all local electrodes. b’) Calculation of local burst synchrony coefficient (LBSC). LBSC is calculated by quantifying the percent change between the mutual information of local and distant electrodes. b’’) LBSC shows enriched local synchrony across all DIVs.

We confirmed the firing behavior of wild-type networks. As expected, number of active electrodes and the mean firing rate increased with time (i.e. network maturation) as seen by a temporal increase in the number of active electrodes and the mean firing rate. We note the number of active electrodes peak and mean firing rate begins to consistently increase by DIV17 (figure 1b,c). Further, we note three established measurements of synchrony (mutual information, entropy and STTC) show evidence of increasing synchrony from DIVs 11-15 and then direction of synchronicity changes no longer correlate between measures from DIVs 17-21 (extended data fgigure 1). Thus, we consider activity patterns separated into evenly sized time period bins that we denote as “Early maturation” and “More Mature”, and show distinct firing rates and number of activity electrodes between these two time periods (figure 1b’,1c’). Next, we tested whether there was increased synchrony among local electrodes in wild-type networks. To this end, we considered two previously described distinct measures of synchrony – Mutual Information and STTC (Gelfman 2018). Mutual Information considers how frequently electrodes burst together, while STTC looks at the overall pairwise correlation of spike-trains using a moving time window. Importantly, Mutual Information and STTC do not necessarily increase or decrease together. Indeed, wild-type networks showed distinct synchrony patterns in developing cultured networks using these existing measures of synchrony. Mutual information increased with time, whereas STTC increased until DIV17 and then plateaued (extended data figure 1).

Similarly, aberrant synchrony *in vitro* in the context of disease modeling or drug perturbation may present with distinct measures of synchrony showing opposing effects, such as an increase in STTC and an increase in Mutual Information in the same plate, despite an increase in Mutual Information being indicative of decreased synchrony (extended data table 1). Such findings are consistent with prior studies that have shown there is significant variation in the direction of synchrony phenotypes when networks were perturbed with addition of GABA and AMPA modulators (Bader 2017, Eisenman 2015). Thus, we aimed to generate two distinct topology-based synchrony measures – the Local Synchrony Coefficient (LSC) and the Local Bursting Synchrony Coefficient (LBSC) in order to capture more potential patterns of aberrant synchrony than one metric alone.

Both the LSC and LBSC examine the relative strength of local networks, with positive values indicative of nearby neurons being more synchronized than more spatially distant neurons. The LSC is derived from STTC, a measure of the similarity of firing patterns overall, while the LBSC is derived from MI, which focuses on whether or not bursting activity is correlated (figure 2a,b). Thus, the LSC and LBSC used two distinct approaches to quantify the relationship between synchrony and distance *in vitro*. Considering both the entire spike train (LSC) and bursting patterns (LBSC), we were able to identify enriched local synchrony in wild-type culture networks, with values greater than 0 at all time points for both metrics (figure 2 c,d).

**Figure 2:**
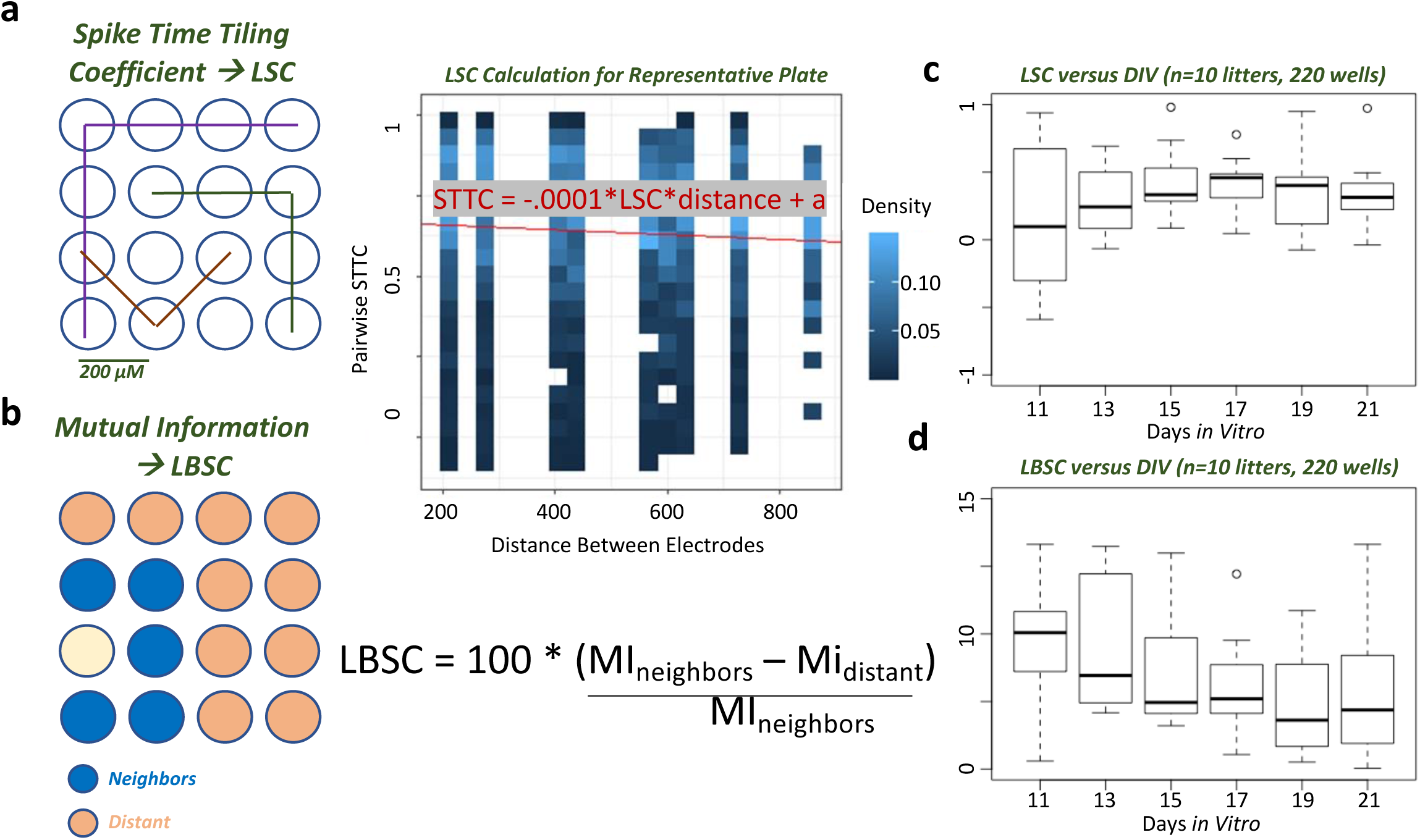
Aberrant Topology in Neuronal Networks Treated with Subtype-specific Activity Modulators. a) Description of experimental approach for early perturbation of activity. b) Chronic dosing with MgCl increases synchrony, as quantified by existing metrics. MgCl dosing leads to an increase in MI across all time points and an increase in STTC for early time points (DIVs 11,13,15). c) Aberrant topology identified in MgCL and Muscimol treated networks is complementary to existing synchrony metrics. MgCl shows aberrant bursting topology concordant with disrupted MI, but no difference in LSC at early or late time points. Muscimol treatment results in aberrant local topology without identifiable synchrony phenotypes using existing synchrony metrics.

We found the developmental timeline of enriched local synchrony varies depending on topological approach. For example, the LSC does not show clear enrichment of local networks at DIV11, while there is evidence of increased local synchrony using LBSC. While both measures elucidate enriched local synchrony, it must be noted that local topology is not apparent in all plates at all later time points, with the presence of negative LSC and LBSC values at DIV21 (figure 2 c,d). Thus, identification of topological phenotypes may be more robust during certain developmental time windows *in vitro*.

### Network topology disrupted by neuronal subtype activity modulators

We next considered to what extent we could perturb network topology and development by modulating the activity of certain neuronal subtypes. Specifically, we examined whether chronic perturbation of cortical networks beginning at DIV3 had longitudinal or long-lasting effects on network activity. We tested compounds known to perturb glutamatergic activity via NMDA receptors (MgCl_2_) or GABAergic activity using the potent and highly selective GABA_A_ agonist, Muscimol (figure 3a), processes known to have significant roles in neurodevelopment (Wu 2015, Wang 2009, Luhmann 2016, Ojeda 2019).

**Figure 3:**
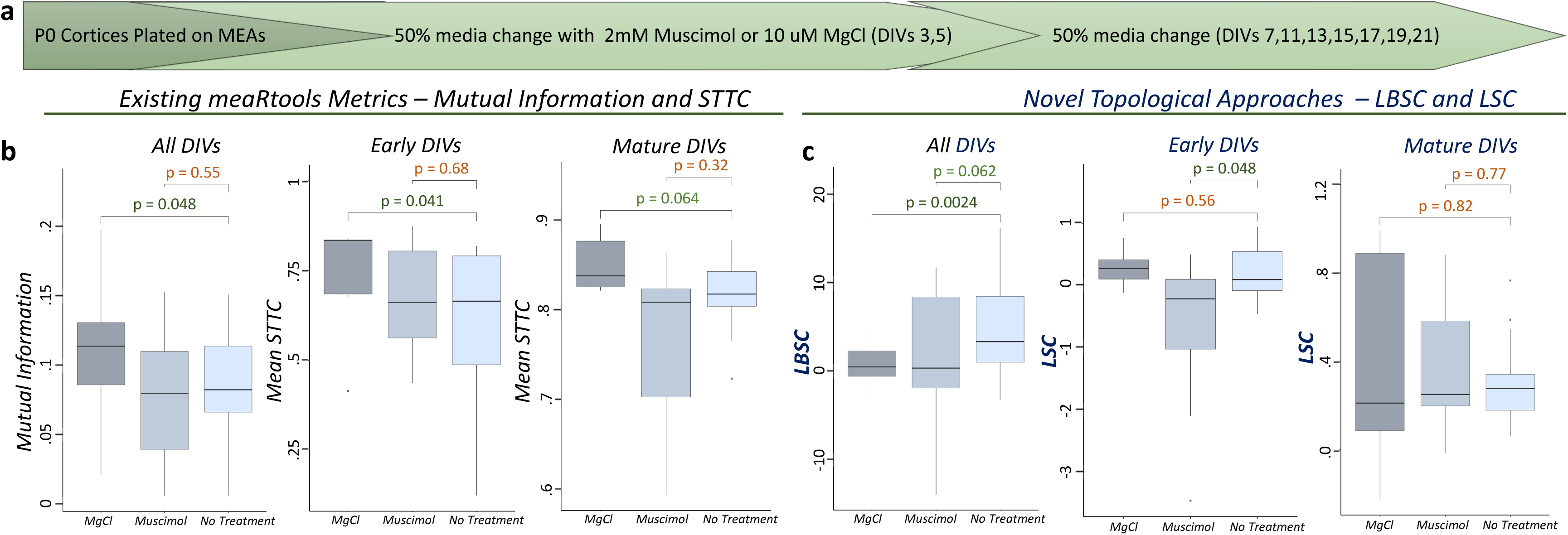
Topological Measures Identify Novel, Complimentary Phenotypes in Mouse Models of Neurodevelopmental Disorders. a) In *Stxbp1^+/−^* mice, LBSC and Mutual Information show significant and trending significant differences, respectively. b) In *Grin2a^S644G/S644G^*, significant differences in Mutual Information does not result in aberrant bursting topology. c) In *Hnrnpu*^+/−^ mice, disruption of local synchrony despite no difference in mean STTC (n=6 plates, 144 wells). Across three models, incorporating topology allows for identification of phenotypes distinct from existing synchrony measures.

We assessed whether treatment of networks during early development (DIV3 and DIV5) with either MgCl or Muscimol resulted in aberrant topology at more mature time points. Importantly, media was replenished every other day, leading to minimal impact of active drug at later time points. We hypothesized modulation of network activity during early development would lead to modest phenotypes later in development, driven by developmental compensation muting the effect of acute drug treatment. While early MgCl addition led to significant synchrony phenotypes using the LBSC measure showing decreased enrichment of synchronicity of neighboring electrodes and Mutual Information showing increased global synchrony (figure 3b,c). Muscimol addition resulted in a consistent and similar ablation of network topology at earlier time points, without a corresponding phenotype in global synchrony, as measured by STTC (figure 3b,c). Thus, spatial analyses of synchrony are capable of identifying phenotypes both concordant and discordant with existing features, suggesting such approaches are complementary to existing algorithms in identifying aberrant synchrony driven by perturbation of activity-dependent processes and may additionally increase sensitivity to identify synchrony phenotypes driven by early developmental disruption.

Existing measures of synchrony (Entropy, MI, STTC) are consistent in their behavior from DIV11-15, but then begin to diverge as the network further matures (see extended data figure 1). Thus, we examined spatial synchrony at DIV11-15 and at more mature time points (DIV17-21) following activity modulation. At more mature time points, evidence of enriched local synchrony in Muscimol treated wells re-emerges. Early ablation of topology in Muscimol-treated wells may be impacted by low levels of Muscimol remaining in culture. Alternatively, networks may recover during development or sensitivity to topological phenotypes may be limited in mature networks by high overall levels of synchrony.

### LBSC and LSC identify novel phenotypes in models of neurodevelopmental disease

We obtained and characterized the seizure activity via video-EEG of a constitutive *Stxbp1^+/−^* mouse model generated as part of the Knockout Mouse Phenotyping program (KOMP). Similar to other genetic mouse models of STXBP1 haploinsufficiency (Kovačević 2018, Chen 2020), the KOMP *Stxbp1*^+/−^ mice displayed a preponderance of absence seizure-like spike-and-wave discharges with increased discharges during dark periods (extended data figure 4). While there was no evidence of spontaneous seizures in *Hnrnpu^+/−^*mice, they exhibit reduced seizure thresholds (Dugger 2020). *Grin2a*^S644G^ heterozygotes and homozygotes showed resistance to electrically induced seizures and lethal tonic-clonic seizures, respectively (Amador 2020).

Next, we asked whether or not topological analyses can identify spatial synchrony phenotypes in addition to existing synchrony measures. To this end, we examined three distinct mouse models of neurodevelopmental disease. Loss of function mutations in *STXBP1* (Saitsu 2008) and *HNRNPU* (Epi4K Consortium 2012, Yates 2017) result in severe early onset epileptic encephalopathies. Similarly, a patient specific gain-of-function mutation (S644G) in the NMDA receptor *GRIN2A* result in a severe development epileptic encephalopathy.

**Figure 4:**
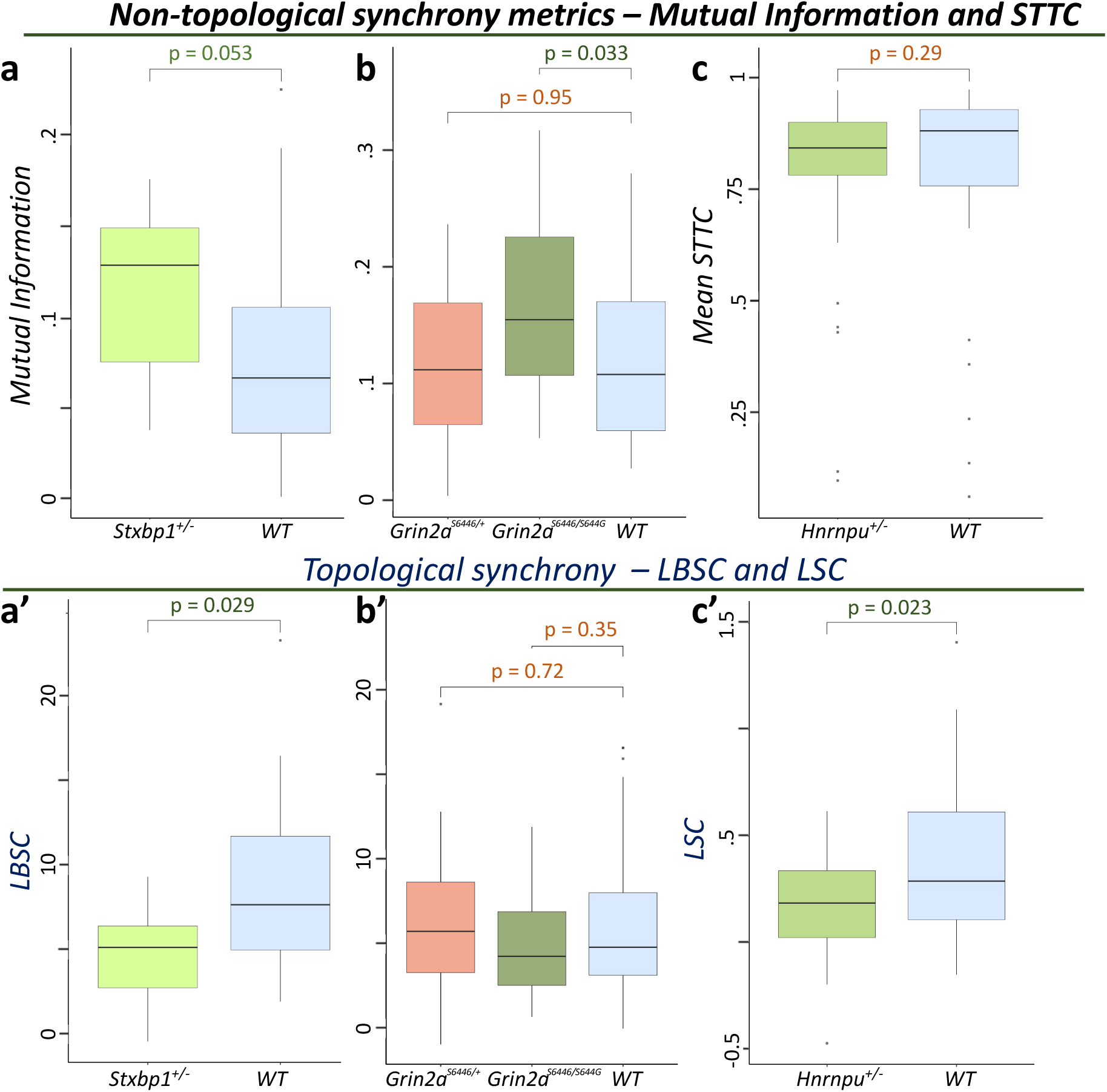
Local Clustering Coefficient (LCC) Identifies Novel Phenotype in Grin2a^S644G/S644G^ Mice. a) LCC is generated by considering all electrode pairs with an STTC >0.5 as ‘connected.’ Local clustering coefficient for all electrodes is equal to the proportion of local electrodes that are connected. b) Synchrony differences in homozygous mice at DIV11 identified in homozygous, but not heterozygous, mice (n = 5 plates, 14-16 wells per plate). c) LCC identifies aberrant topology in both *Grin2a^+/S644G^* and *Grin2a^S644G/S644G^* mice at DIV11. All conditions become hyper-connected by DIV15.

Cultured neuronal networks from a genetic mouse model of *Grin2a*S644G result in activity and synchrony phenotypes that depend on copy number of the S644G variant (Amador 2020). We examined cultured cortical networks from *Stxbp1^+/−^* and *Hnrnpu^+/−^* mice, and heterozygous (*Grin2a*^S644G/+^) or homozygous (*Grin2a*^S644G/S644G^) mice (see Materials and Methods). *Stxbp1*^-/-^ and *Hnrnpu*^-/-^ networks were not generated because homozygotes exhibit perinatal and prenatal lethality, respectively (Verhage 2000, Dugger 2020).

*Hnrnpu*^+/−^ and *Stxbp1*^+/−^ cortical networks were first assessed for an array of spiking and bursting network features (Gelfman 2018). For *Stxbp1*^+/−^ networks, we observed consistent disruption for several features, including all litters displaying an increase in the number of spikes in bursts (extended data table 1), which suggests a robust phenotype that may be suitable for testing therapeutics. Confoundingly, certain features in both *Hnrnpu*^+/−^ and *Stxbp1*^+/−^ cortical networks showed highly significant increases in aggregate data from one litter as well as a discordant decrease in aggregate data from other litters. Importantly, such discordance is especially pronounced in networks from *Hnrnpu*^+/−^ mice (discordant results in 9 out of 13 activity features, extended data table 1) suggesting MEA may be limited in their specificity for mouse models with more mild *in vivo* phenotypes such as Hnrnpu.

We then analyzed *Hnrnpu*^+/−^ and *Stxbp1*^+/−^ cortical networks, as well as previously described *Grin2a* S644G heterozygous and homozygous cultures, using the LSC and LBSC metrics to determine whether such topological measures can improve upon the conventional measures of synchrony, STTC and MI. In *Hnrnpu*^+/−^ mice, LSC shows aberrant topology in mutant networks independent of mean STTC (figure 4c,4c’). Using the LBSC, *Stxbp1*^+/−^ networks exhibit aberrant topology as well as a corresponding increase in Mutual Information, suggesting there are synchrony phenotypes in Stxbp1 robust enough to present across a variety of algorithmic approaches (figure 4a,a’). Importantly, a significant increase in Mutual Information alone does not necessarily lead to an aberrant topological bursting phenotype, as there is no significant difference in LBSC in *Grin2a**^S^***^644G/S644G^ mice (figure 3b,b’). Thus, topological approaches such as LSC and LBSC identify spatial synchrony phenotypes where the relationship between synchrony and distance is disrupted in certain mouse models of neurodevelopmental disorders. Such spatial synchrony measures are complementary to distant agnostic synchrony measures such as STTC. Importantly, they are capable of identifying synchrony phenotypes both in models with significant disruption, such as *Stxbp1*, and those that have more muted *in vivo* phenotypes, as exemplified by the *Hnrnpu*^+/−^ mouse model.

### Local Clustering Coefficient identifies early disruption in *Grin2a***^+^**^/S644G^ mice

Once we established incorporating distance between electrodes compliments and extends existing synchrony measures, we leveraged graph theory to generate an adaptation of the commonly used clustering coefficient. We used this additional topological measure, the Local Clustering Coefficient (LCC) (figure 5a) to assess whether aberrant local topology could be identified in cultured networks from *Grin2a*^S644G^ heterozygous mice. Evidence of enriched local synchrony shown by the LSC (figure 2) motivated a focused computational tool specifically investigating local networks. The LCC similarly looks at pairwise correlations of spike-trains, but only considers neighboring electrodes and designates each electrode pair as synchronized or not based on a simple STTC threshold value of 0.5. Importantly, during network development, it has previously been shown that Grin2a^S644G/+^ networks resemble wild-type when assessed by Mutual Information and STTC, whereas *Grin2a*^S644G/S644G^ networks exhibit clear synchrony differences relative to wild-type (Amador 2020). Using the LCC, we found significant reduction in local clustering Grin2a^S644G/+^ networks without a reduction using global synchrony metrics (figure 5b,c) at DIV11.

**Figure 5:**
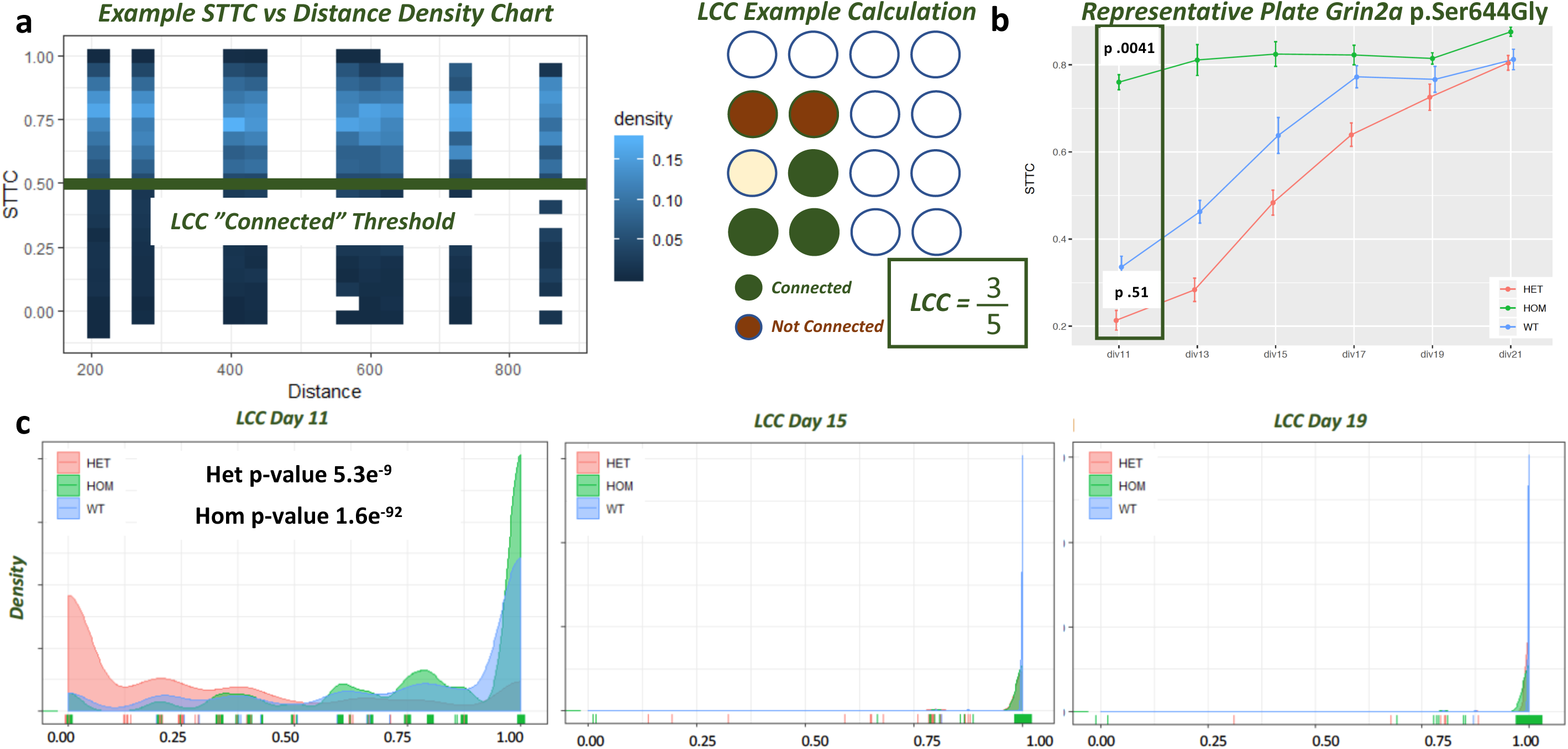
Local Clustering Coefficient (‘LCC’) Identifies Novel Phenotype in Grin2a^p.Ser644Gly/+^ Mice. a) LCC is generated by considering all electrode pairs with an STTC >0.5 as ‘connected.’ Local clustering coefficient for all electrodes is equal to the proportion of neighboring electrodes that are connected. b) Synchrony differences in homozygous mice at DIV11 identified in homozygous, but not heterozygous, mice (n = 5 plates, 14-16 wells per plate). c) LCC identifies aberrant topology in both *Grin2a^p.Ser644Gly/+^ and Grin2a^p.Ser644Gly/p.Ser644Gly^* mice at DIV11. All conditions become hyper-connected by DIV15.

The LCC also allows for visualization of how network connectivity evolves over time. In a representative plate, a wide spectrum of LCCs is seen at DIV11, however, nearly all electrode pairs surpass the ‘connected’ threshold by DIV15 for all conditions (figure 5c). Such hyper synchronization can also be seen as the mean STTC for wild-type and mutant mice begin to converge by DIV21 (figure 5b). The rapid saturation of LCC suggests the feature is best suited to identify early developmental phenotypes and is unlikely to be sensitive to aberrant topology in more mature networks.

## Discussion

This study focuses on first characterizing the developmental trajectory of the relationship between proximal (local) versus distal (distant) synchrony in murine cortical neuronal networks. Then, we utilize broadly acting compounds to show the enrichment of local synchrony can be perturbed through altered activity. Further, we investigate whether such topological measures increase the dynamic range of MEAs to identify novel phenotypes independent of existing measures of synchrony. The topological measurements presented in this work were generated with the intention to both examine the presence of increased synchrony of local electrodes in a medium-throughput MEA system and to determine whether such measures are capable of identifying disease-relevant phenotypes. We leveraged the existence of previously described measures of synchrony and adapted aspects of graph theory measures to assess the effects on synchrony. We used drug perturbation of primary neuronal networks to test our hypotheses and refine our methods, and then tested the newly developed methods on neuronal networks from three genetic mouse models of neurodevelopmental disease.

By identifying the presence of enriched local synchrony in 48-well MEA plates, we established that systems with a limited number of electrodes can be utilized for a wide array of topological analyses, including, but not limited to, those described within this work. Consistent with the divergent developmental patterns of STTC and Mutual Information, the pattern of enrichment of local synchrony using the LSC and LBSC suggest the measures are identifying distinct topological features. Neuronal networks are incredibly complex, as are the molecular mechanisms that generate specific topological patterns. We do not address the molecular underpinnings driving local topology and further molecular biology experiments are required to better understand the drivers of LSC and LBSC.

While we do not address specific molecular mechanisms, we do show that broad activity-dependent processes may drive aberrant topology *in vitro*. We show that early perturbation of both Glutamatergic and GABAergic activity ablates elements of local topology. Further, we note that all significant LSC and LBSC differences identified here in mouse models of neurodevelopmental disorders show a reduction in enrichment of local synchrony. One hypothesis is that the models we studied involve hypersynchrony *in vivo* and hypersynchrony may lead to reduced enrichment of local synchrony *in vitro*, which is consistent with an overall trend of reduced LSC and LBSC at later time points of wild-type cultures (figure 2).

The three measures described – LSC, LBSC, LCC – enable the identification of screenable phenotypes for testing potential therapeutics. Using these measures, we were able to identify novel topological phenotypes in three distinct disease models, which are complementary to existing synchrony measures. Of particular interest, we note we identified robust synchrony phenotypes in *Stxbp1^+/−^* mice *in vivo* and *in vitro* with both global and topological approaches to measuring synchrony, suggesting such a model may be particularly useful for therapeutic development using MEAs, with evaluation initially based on restoring global and topological synchrony *in vitro*, followed by validation *in vivo*.

In general, topological phenotypes are sensitive to the functional maturity of a given neuronal network. For this reason, wild-type networks show no evidence of increased local synchrony at certain DIVs, likely due to hyper-synchronization. Further, in some cases aberrant topology may only be identified with one of the features described here. Given the significant plate to plate variability across conventional features in MEAs, singular topological phenotypes may be artifacts in certain circumstances. Further, well-to-well and plate-to-plate variability and high levels of synchrony may limit the capacity to identify aberrant topology which is an inherent limitation of the MEA platform. These drawbacks necessitate the examination of a large number so replicates thereby reduces the throughput of MEA for therapeutic screens. While an optimal experimental design for identifying MEA synchrony phenotypes *in vitro* remains challenging, we would suggest at least 5 plates of recordings, as that number was utilized to identify topological synchrony phenotypes within this work. Additionally, any therapeutic screening we would suggest requiring at least two global or topological synchrony phenotypes to be reversed for any compound to be considered interesting given the inherent variability of MEA networks and synchrony algorithms. The age of networks to be examined should be determined *a priori* to avoid selection bias. One potential approach, as we did within this manuscript, is determine time periods to examine based on number of active electrodes and the plateauing of certain synchrony metrics, such as STTC.

In conclusion, we have generated a set of topological measures that complement existing measures in mouse models we suggest are ideally suited for therapeutic development using MEAs, such as *Stxbp1*, while improving upon existing measures of synchrony to identify novel phenotypes in *ex vivo* neuronal networks from mouse models of neuodevelopemtal disorders with more muted *in vivo* and *in vitro* phenotypes, such as *Hnrnpu* and *Grin2a*.

## Author Contributions

AR – designed research, performed research, analyzed data and wrote the paper. SD designed research and performed research. SC, SP and DK performed research. WF, DG and MB designed research.

## Acknowledgments

We thank Sahar Gelfman and Michael Beaumont for initial discussions on topological tools for MEAs, and for consideration of approaches that laid the groundwork for the algorithms described here.

## Conflict of interest

D.B.G. is a founder of and holds equity in Praxis and Actio Bioscience. All other authors declare no competing financial interests.

---

**Extended Data Figure 1:**
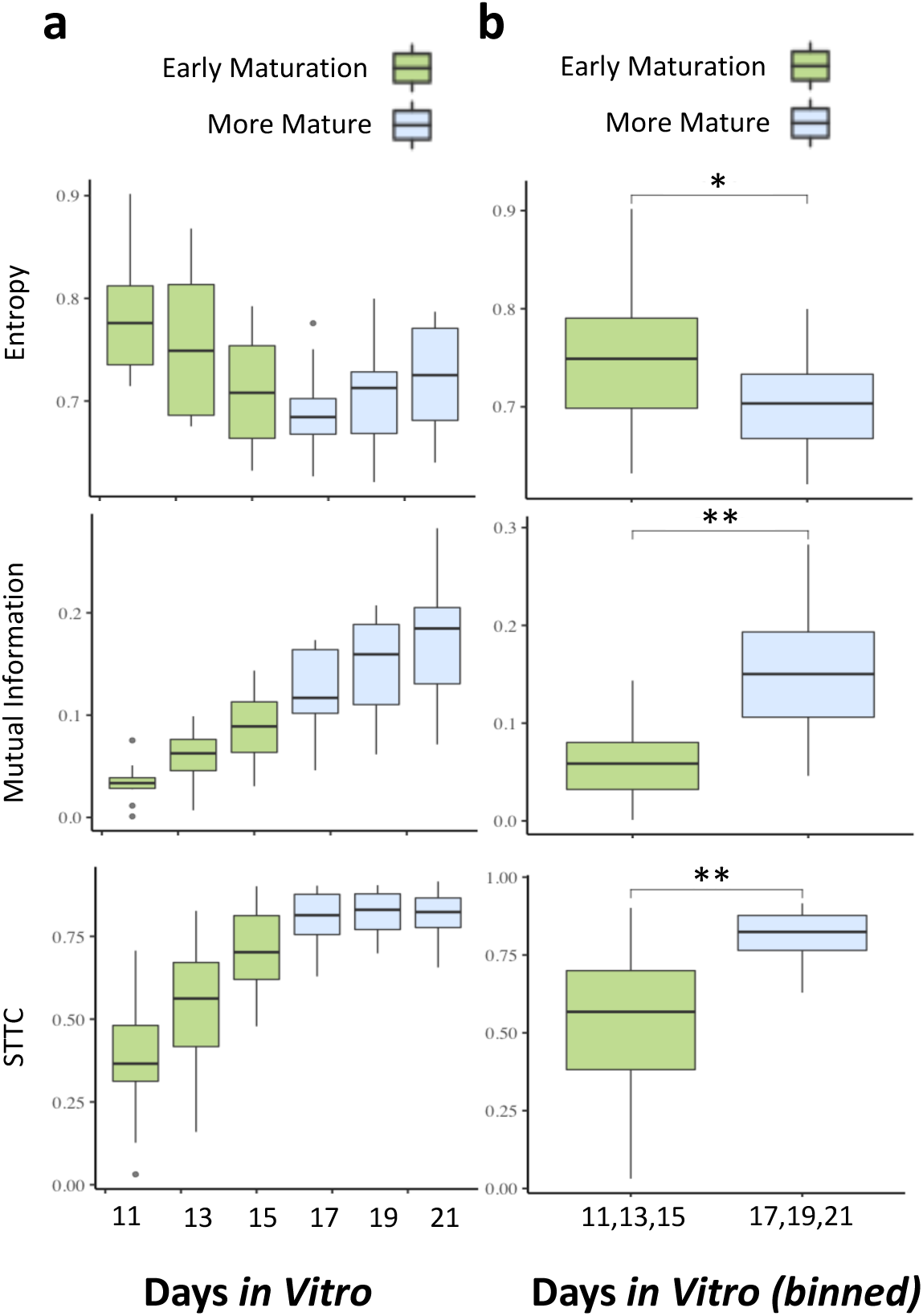
Synchrony features behave differently over time. We explored the relationship between a subset of synchrony phenotypes from meaRtools. Box plots for n = 10 litters, 220 wells with each box corresponding to a specific DIV (a) or set of DIVs (b). Synchrony phenotypes consistently increase during early maturation (Lower entropy = increased synchrony). However, synchrony phenotypes have incongruous patterns in more mature time points. All three synchrony features show differences between early and more mature time periods (* < .05, ** p< .001). Mean Firing Rate corresponds to the mean firing rate per active electrode. STTC = Spike-time tiling coefficient.

**Extended Data Figure 4:**
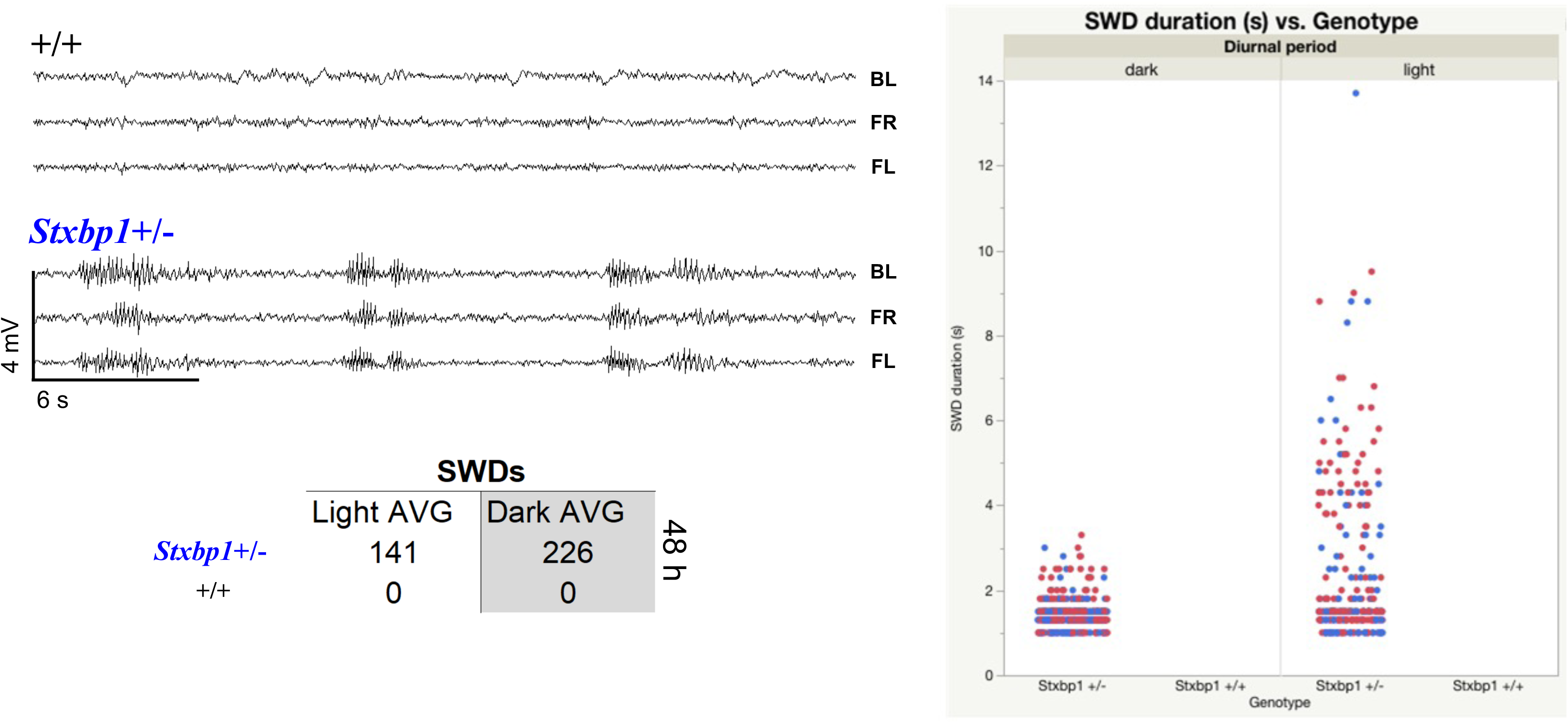
Spike-and-Wave Discharges in *Stxbp1^+/−^* mice*. Stxbp1^+/−^* mice display absence seizure-like EEG phenotypes *in vivo*. Significant number of SWDs in both light and dark, with higher frequency, shorter discharges in the dark period. a) Example traces from electrodes implanted in wild-type and *Stxbp1^+/−^* mice. “BL” = back left. “FR” = front right. “FL”. b) Duration of all identified SWDs in 7 *Stxbp1*^+/−^ mice and 3 wild-type litter mates. c) Average number of SWDs in light and dark periods over 48 hours (n = 7 *Stxbp1*^+/−^, 3 wild-type).

**Extended Data Table 1:** MEA Characterization of Hnrnpu and Stxbp1 haploinsufficiency in primary cortical cultures. Permutation p-values and the direction of change listed across 13 features analyzed using meaRtools software. “nAE” = number of active electrodes. “MFR/nAE” = Mean firing rate per active electrode. “Mean Dur of Bursts” = Mean duration of bursts. “% spikes in NS” = percentage of spikes in networks spikes. “mean spikes in NS” = average number of spikes in a network spike. Plate to plate variability especially apparent in *Hnrnpu*^+/−^ networks, with opposing significant differences in 9 out of 13 features considered.

| Mouse Model |  | *Hnrnpu*<sup>+/-</sup> |  |  |  |  |  | Mouse Model |  | *Stxbp1*<sup>+/-</sup> |  |  |  |
| --- | --- | --- | --- | --- | --- | --- | --- | --- | --- | --- | --- | --- | --- |
| Plate ID |  | 1 | 2 | 3 | 4 | 5 | 6 | Plate ID |  | 1 | 2 | 3 | 4 |
| #wells | WT | 24 | 24 | 24 | 24 | 24 | 24 | #wells | WT | 24 | 24 | 24 | 24 |
|  | Het | 24 | 24 | 24 | 24 | 24 | 24 |  | Het | 24 | 24 | 24 | 24 |
| DIVs | All | DIV11-21 | DIV11-21 | DIV11-21 | DIV11-21 | DIV11-21 | DIV11-21 | DIVs | All | DIV11-21 | DIV11-21 | DIV11-21 | DIV11-21 |
| nAE | perm_pval | 0.16 | 0.01 | 0.01 | 0.21 | 0.01 | 0.01 | nAE | perm_pval | 0.61 | 0.96 | 0.01 | 0.16 |
|  | direction | No Difference | Increase | Decrease | No Difference | increase | Increase |  | direction | No Difference | No Difference | Increase | No Difference |
| MFR/nAE | perm_pval | 0.07 | 0.07 | 0.57 | 0.01 | 0.01 | 0.01 | MFR/nAE | perm_pval | 0.17 | 0.42 | 0.01 | 0.19 |
|  | direction | No Difference | No Difference | No Difference | Increase | increase | Increase |  | direction | No Difference | No Difference | Increase | No Difference |
| MutInfo | perm_pval | 0.74 | 0.02 | 0.16 | 0.01 | 0.86 | 0.01 | MutInfo | perm_pval | 0.03 | 0.01 | 0.01 | 0.25 |
|  | direction | No Difference | Increase | No Difference | Increase | No Difference | Increase |  | direction | Decrease | Increase | Increase | No Difference |
| Entropy | perm_pval | 0.46 | 0.01 | 0.01 | 0.01 | 0.01 | 0.01 | Entropy | perm_pval | 0.01 | 0.01 | 0.01 | 0.10 |
|  | direction | No Difference | Decrease | Increase | Increase | Increase | Increase |  | direction | Decrease | Decrease | Decrease | No Difference |
| STTC | perm_pval | 0.49 | 0.01 | 0.11 | 0.97 | 0.01 | 0.01 | STTC | perm_pval | 0.01 | 0.01 | 0.01 | 0.82 |
|  | direction | No Difference | Increase | No Difference | No Difference | Decrease | Decrease |  | direction | Increase | Increase | Increase | No Difference |
| Number of Bursts | perm_pval | 0.44 | 0.11 | 0.68 | 0.33 | 0.01 | 0.01 | Number of Bursts | perm_pval | 0.01 | 0.01 | 0.23 | 0.01 |
|  | direction | No Difference | No Difference | No Difference | No Difference | increase | Increase |  | direction | Decrease | Decrease | No Difference | Decrease |
| Mean Dur of Bursts | perm_pval | 0.04 | 0.04 | 0.91 | 0.01 | 0.02 | 0.01 | Mean Dur of Bursts | perm_pval | 0.54 | 0.00 | 0.04 | 0.01 |
|  | direction | Decrease | Decrease | No Difference | Increase | Increase | Increase |  | direction | No Difference | Increase | Increase | Increase |
| Mean Frequency in Burst | perm_pval | 0.11 | 0.01 | 0.52 | 0.01 | 0.01 | 0.73 | Mean Frequency in Burst | perm_pval | 0.01 | 0.14 | 0.01 | 0.10 |
|  | direction | No Difference | Increase | No Difference | Decrease | Decrease | No Difference |  | direction | Increase | No Difference | Increase | No Difference |
| Mean Spikes in Bursts | perm_pval | 0.01 | 0.95 | 0.01 | 0.01 | 0.01 | 0.01 | Mean Spikes in Bursts | perm_pval | 0.01 | 0.01 | 0.01 | 0.01 |
|  | direction | Decrease | No Difference | Decrease | Increase | Bidirectional | Increase |  | direction | Increase | Increase | Increase | Increase |
| % spikes in Burst | perm_pval | 0.31 | 0.09 | 0.27 | 0.02 | 0.01 | 0.01 | % spikes in Burst | perm_pval | 0.29 | 0.01 | <.0001 | 0.01 |
|  | direction | No Difference | No Difference | No Difference | Increase | increase | Decrease |  | direction | No Difference | Increase | Increase | Increase |
| # Network spikes | perm_pval | 0.56 | 0.14 | 0.55 | 0.01 | 0.01 | 0.01 | # Network spikes | perm_pval | 0.01 | 0.01 | 0.60 | 0.01 |
|  | direction | No Difference | No Difference | No Difference | Increase | Increase | Increase |  | direction | Decrease | Decrease | No Difference | Decrease |
| % spikes in NS | perm_pval | 0.56 | 0.01 | 0.84 | 0.01 | 0.01 | 0.01 | % spikes in NS | perm_pval | 0.01 | 0.01 | 0.06 | 0.01 |
|  | direction | No Difference | Increase | No Difference | Decrease | Decrease | Decrease |  | direction | Decrease | Decrease | No Difference | Decrease |
| mean spikes in NS | perm_pval | 0.01 | 0.01 | 0.04 | 0.01 | 0.01 | 0.72 | mean spikes in NS | perm_pval | 0.01 | 0.01 | 0.01 | 0.90 |
|  | direction | Decrease | Increase | Decrease | Decrease | Decrease | No Difference |  | direction | Increase | Increase | Increase | No Difference |

